# A Mechanism for Neurofilament Transport Acceleration through Nodes of Ranvier

**DOI:** 10.1101/806786

**Authors:** Maria-Veronica Ciocanel, Peter Jung, Anthony Brown

## Abstract

Neurofilaments are abundant space-filling cytoskeletal polymers in axons that are transported along microtubule tracks. Neurofilament transport is accelerated at nodes of Ranvier, where axons are locally constricted. Strikingly, these constrictions are accompanied by a sharp decrease in neurofilament number but no decrease in microtubule number, bringing neurofilaments closer to their microtubule tracks. We hypothesize this leads to an increase in the proportion of the time that the filaments spend moving and that this can explain the local acceleration. To test this, we developed a stochastic model of neurofilament transport that tracks their number, kinetic state and proximity to nearby microtubules in space and time. The model assumes that the probability of a neurofilament moving is dependent on its distance from the nearest available microtubule track. Taking into account experimentally reported numbers and densities for neurofilaments and microtubules in nodes and internodes, we show that the model is sufficient to explain the local acceleration of neurofilaments across nodes of Ranvier. This suggests that proximity to microtubule tracks may be a key regulator of neurofilament transport in axons, which has implications for the mechanism of neurofilament accumulation in development and disease.

## Introduction

Nerve cells extend long cellular processes called axons and dendrites which form electrical connections with other cells throughout the body, thereby establishing the wiring pattern of the nervous system. Communication along these cellular conduits is achieved by the propagation of action potentials, which are waves of membrane depolarization commonly referred to as nerve impulses. Two fundamental mechanisms by which animals can increase the rate of propagation of nerve impulses along axons are to increase axon diameter or to insulate the axon by myelination (Waxman, 1980; Hartline and Colman, 2007). Myelination is a tight spiral wrapping of an axon by a sheet-like extension of a myelinating glial cell. The myelin sheath along a single axon is arranged in contiguous segments called internodes, each formed by a single myelinating glial cell (Schwann cells in the peripheral nervous system, oligodendrocytes in the central nervous system). Each myelinated internode is separated from the next by a short gap of bare axon known as a node of Ranvier, where ion channels that are responsible for initiation and propagation of the nerve impulse (action potential) are concentrated. By clustering the ion channels at nodes and insulating the axon between nodes, myelinated axons are able to propagate nerve impulses in a saltatory manner in which the depolarization at each node spreads rapidly to the next within the myelinated internode (Stampfli, 1954; Salzer, 2003).

For many decades it has been known that myelinated axons are constricted locally and abruptly at nodes of Ranvier (Hess and Young, 1952; Berthold, 1978) and that the extent of constriction scales with the internodal axon diameter (Rydmark, 1981; Sward et al, 1995). Using computational modeling, we and others have shown that these constrictions increase the efficiency of saltatory nerve conduction by decreasing nodal capacitance, thereby reducing the internodal caliber required to achieve a given target conduction velocity by as much as 3-fold (Halter and Clark, 1993; Johnson et al, 2015). We also found that there is an optimum theoretical extent of nodal constriction for any given internodal caliber, and that this matches the extent of constriction observed in animals (Johnson et al, 2015). Thus, nodal constrictions appear to be an evolutionary adaptation that confers significant spatial and metabolic efficiency on myelinated axons.

Because of their long length, axons are critically dependent on the intracellular transport of organelles and macromolecules for their growth and survival. This movement is called axonal transport (Brown, 2016). In addition to their electrophysiological significance described above, nodes of Ranvier also have important implications for the mechanisms of axonal transport because they represent potential bottlenecks for the movement of axonally transported cargoes. One of the most abundant cargoes in the axon are neurofilaments, which are long flexible space-filling protein polymers that function to expand axon caliber (Hoffman, 1995). Neurofilaments move along microtubule tracks in a rapid intermittent and bidirectional manner, alternating between short bouts of rapid anterograde or retrograde movement and long bouts of pausing (Wang et al, 2000; Brown, 2014).

Electron microscopic studies of axons have shown that the number of neurofilaments declines at nodes of Ranvier by as much as 10-fold in the largest axons whereas the number of microtubules does not decline (Tsukita and Ishikawa, 1981; Reles and Friede, 1991, Price et al, 1990; Price et al, 1993; Hsieh et al, 1994). This suggests that most microtubules course through the node from one internode to the next whereas most neurofilaments do not. Recently, we showed that neurofilaments navigate these axonal constrictions by accelerating locally, analogous to the increase in the current where a river narrows its banks (Walker et al, 2019). However, the mechanism of this local acceleration remains unclear. One possible explanation is that neurofilament transport is regulated in part by the proximity of the neurofilaments to their microtubule tracks. Neurofilaments in internodes greatly outnumber microtubules and thus many neurofilaments are not adjacent to a microtubule. In contrast, the ratio of neurofilaments to microtubules is much lower in nodes and thus the average distance between neurofilaments and the nearest microtubule is less at these sites. Here we use computational modeling to test whether this difference in organization is sufficient to explain the local acceleration of neurofilaments in nodes.

## Model

### General description

We model neurofilament movement using an extension of a previous cargo-based model (Jung and Brown, 2009; Li et al, 2012), where each neurofilament moves bi-directionally along the axon cycling between six kinetic states. In the “on-track” states *a*, *a*_0_, *r*, *r*_0_, neurofilaments are associated with microtubule tracks and exhibit movement in anterograde and retrograde directions (states *a* and *r*), interrupted by brief pauses in the resting states *a*_0_ and *r*_0_. During the brief pauses, neurofilaments can reverse direction. The dwell times in these pausing states are of the order of seconds to minutes, resulting in a stop-and-go motion of neurofilaments cycling stochastically between the pausing and moving states (Brown et al, 2005). Using computational modeling, we demonstrated that these four states capture the movement of neurofilaments on short time scales of the order of seconds and minutes but not on longer time scales of the order of hours and days (Brown et al, 2005). To account for this discrepancy, we proposed that neurofilaments in the on-track pausing states *a*_0_ and *r*_0_ can switch to corresponding “off-track” prolonged pausing states *a_p_* and *r_p_*, in which we envisioned that neurofilaments were temporarily disengaged from their microtubule tracks, as if parked on the side of the road. The transitions between the on-track and off-track pausing states were dictated by the rate constants γ*_on_* and γ*_off_*. Subsequently, we confirmed the existence of these distinct pausing states experimentally in cultured neurons and peripheral nerve axons ex vivo using a fluorescence photoactivation pulse-escape technique (Trivedi et al, 2007; Monsma et al, 2014; Li et al, 2014; Walker et al, 2019). The resulting six-state model is depicted in Figure 1.

**Figure 1.**
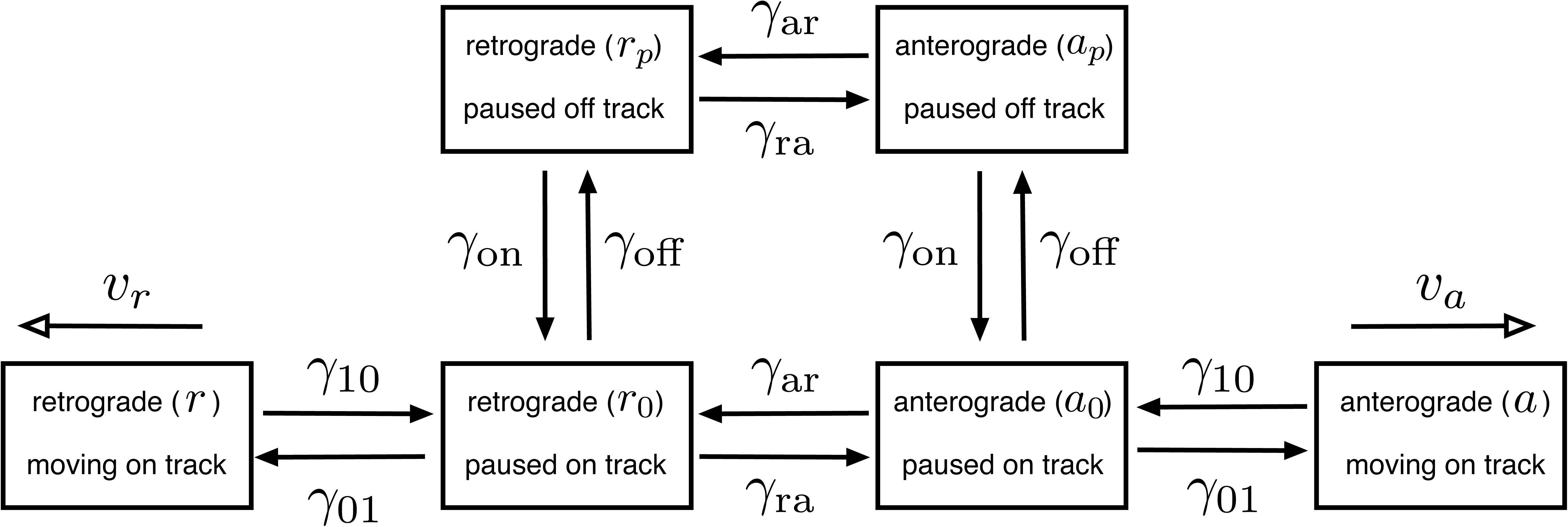
Diagram of the six-state kinetic model of neurofilament transport. There are four on-track states (*a*, *a*_0_, *r*, *r*_0_) and two off-track states (*a_p_*, *r_p_*). On-track neurofilaments move along microtubules in an anterograde or retrograde direction (states *a* and *r*, respectively) with velocities *v_a_* and *v_r_*. The anterograde movements are powered by kinesin motors and the retrograde movements by dynein motors. While in the on-track moving states, the filaments can switch to on-track pausing states *a*_0_ and *r*_0_, governed by the rate γ_10_. When in the on-track pausing states, the filaments can either switch back to their respective on-track moving states, governed by the rate γ_01_, or they can switch to the corresponding anterograde and retrograde off-track pausing states governed by the rates *a _p_* or *r_p_*. Cycling between the on and off-track pausing states is γ*_off_* and γ*_on_*. Reversals in direction can happen in all pausing states, governed by the reversal rate constants, γ*_ar_* and γ*_ra_*. Adapted from Li et al, 2012.

This above model has been instrumental in predicting the kinetics of neurofilament transport in vivo. For example, the model revealed that a pulse of radiolabeled neurofilaments forms a Gaussian wave which moves and spreads at rates consistent with the published experimental data, demonstrating that the rapid intermittent movement of neurofilaments observed in cultured neurons can explain the population behavior of these polymers in animals (Brown et al, 2005; Jung and Brown, 2009). The model has also allowed us to gain insight into the kinetics of neurofilament transport in the optic nerve (Li et al, 2012) and the local regulation of neurofilament transport by myelinating glia (Monsma et al, 2014). However, a significant shortcoming of this model is that it offers no insight into the mechanistic basis for any differences in kinetic behavior and does not relate neurofilament content to axon caliber. Here, we address this shortcoming by incorporating features of the cytoskeletal organization of the axon into the six-state kinetic model, allowing us to explore the influence of the proximity of neurofilaments to their microtubule tracks on the transport kinetics.

### Neurofilament and microtubule organization

The microtubules and neurofilaments are considered to be linear structures arranged in a parallel array coaligned with the long axis of the axon. The microtubules extend the entire length of the axonal domain and form stable tracks for neurofilament transport whereas the neurofilaments are much shorter, with a range of lengths discretized to the nearest integer μ*m* (see Figure 2A). The neurofilaments move forwards and backwards along the microtubules and can diffuse laterally, i.e. in the radial dimension of the axon, when they are not moving.

**Figure 2.**
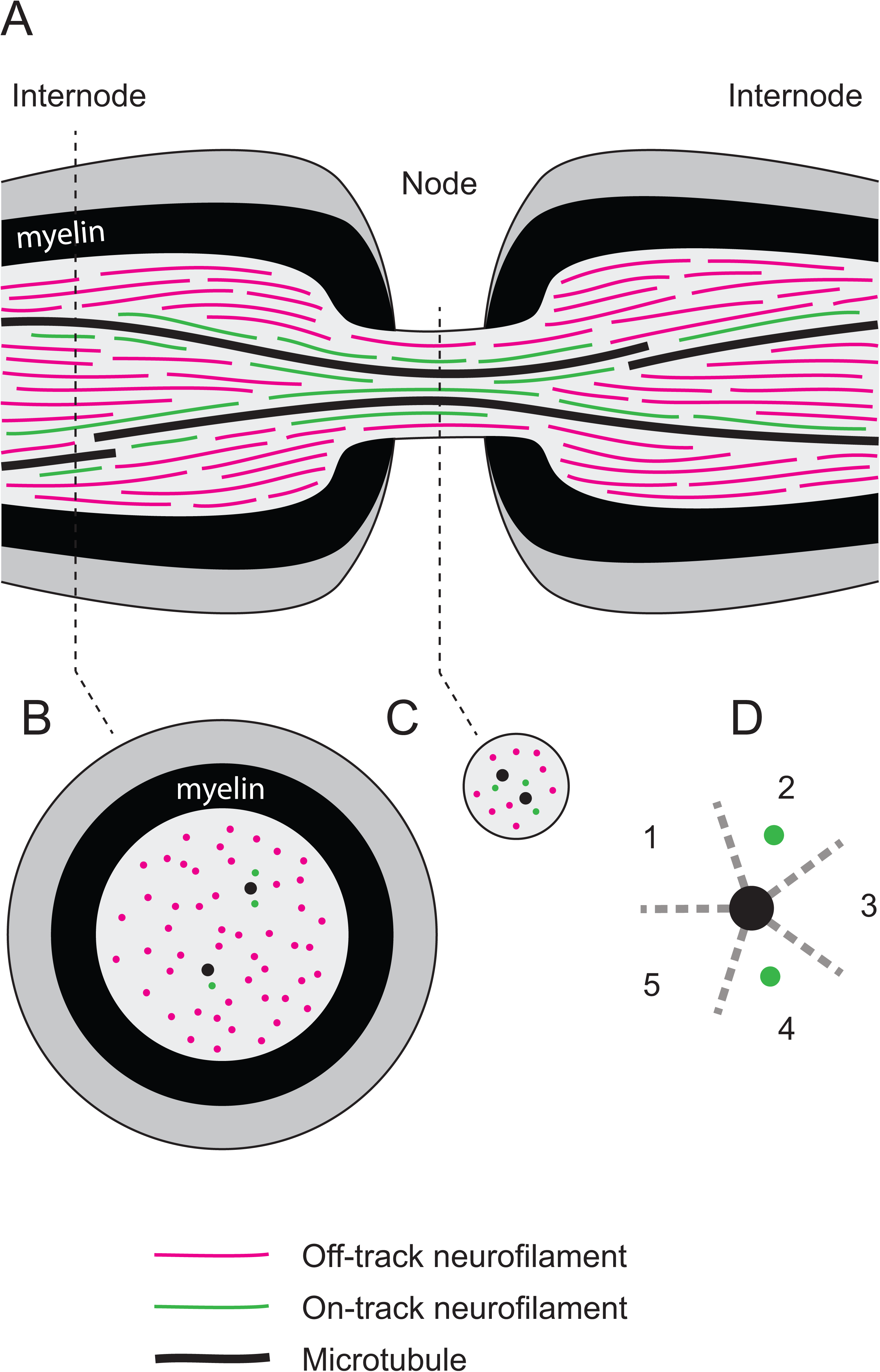
Schematic of a node of Ranvier along a myelinated axon. (A) Longitudinal section through the node and flanking internodes. Neurofilaments switch between on-track (green) and off-track (magenta) states. On-track neurofilaments are engaged with a microtubule track (black) and move along that track in a rapid, intermittent and bidirectional manner. Off-track neurofilaments are disengaged from their microtubule tracks and may get pushed aside, pausing for prolonged periods before re-engaging and resuming movement. To move on track, off-track neurofilaments must diffuse laterally until they encounter a microtubule. (B) Cross-sectional view of the internode. (C) Cross-sectional view of the node. Note that most microtubules run continuously through the node from one internode to the next whereas most neurofilaments terminate on either side of the node, resulting in far fewer neurofilaments in the node than in the internodes. Thus, the average distance between neurofilaments and microtubules is less in the node than in the internodes. (D) View of one microtubule in cross-section (black) with two on-track neurofilaments (green). Due to spatial constraints, each microtubule track is considered to accommodate up to five “lanes” of traffic (numbered 1-5 and separated by dashed grey lines), i.e. a maximum of five neurofilaments at one time (Lai et al, 2018).

Each microtubule has thirteen protofilaments and therefore is theoretically capable of supporting thirteen lanes of traffic. In reality, however, steric considerations are expected to limit the number of neurofilaments that can engage simultaneously at the same location along a microtubule to a maximum of *p* = 5 lanes (see Figure 1 in Lai et al, 2018 and Figure 2D). The microtubules are assumed to be distributed uniformly throughout the axon, meaning that each microtubule has an equal probability of being at any location within the radial dimension of the axon. In such a case, the average distance of a microtubule to its nearest neighbor is given by *d*_0_ = 1/(2ρ^1/2^) (Hertz, 1909) where ρ*_MT_* denotes the density of microtubules, i.e. the number of microtubules per μ*m*^2^. The microtubule and neurofilament densities depend on the specific type of axon and, in general, on their cross-sectional area and can be obtained from published morphometric studies. For axons of the mouse sciatic nerve, which are modeled in the present study, the internodal microtubule density is about 10/μ*m*^2^ and almost independent of axon caliber (Reles and Friede, 1991), resulting in an average nearest distance between microtubules of about 158nm.

### The rate of finding a microtubule track

Inspection of electron micrographs of neurofilament-rich axons reveals that the neurofilaments greatly outnumber the microtubules and that consequently some neurofilaments are adjacent to a microtubule and others are not (Friede and Samorajski, 1970; Price et al, 1988; Reles and Friede, 1991). Thus the probability of moving cannot be the same for all neurofilaments, and neurofilaments that are not next to a microtubule must move laterally in order to become on track. In our six-state model, the rate at which an off-track neurofilament moves on track is governed by the rate constant γ*_on_* (Figure 1). In our new model we consider this process to require a diffusive search in the radial dimension of the axon and thus γ*_on_* becomes an emergent parameter that depends on the average distance between the neurofilaments and their tracks, i.e. the relative cross-sectional densities of these cytoskeletal elements. To define the rate of this radial diffusion, we calculate the mean first passage time for a neurofilament to reach the average distance to the nearest microtubule once it detached, which is given by *T* = *r*^2^/4*D* (Redner, 2001), where *D* denotes the diffusion coefficient of a neurofilament in the axonal cytoplasm and *r* is the average distance to the nearest microtubule (nearest neighbor distance averaged over all angles), defined by 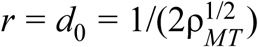. This results in the estimate of

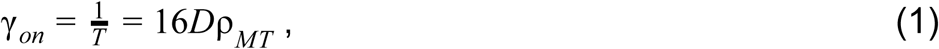

which could be considered the maximum on rate assuming that when a neurofilament meets a microtubule it will always bind to it. While this approximation neglects the geometric effects of microtubule and neurofilament size, it does capture the dependence of the on rate on the microtubule density. When many neurofilaments are present and each microtubule can be engaged with only a finite number of neurofilaments *p*, the binding probability for any given neurofilament is reduced. Denoting by *N_on_* the number of neurofilaments engaged with the microtubule at a specific location along the axon, i.e. the number of neurofilaments in the on-track moving (*a*, *r*) and on-track pausing (*a*_0_, *r*_0_) states, then the number of *available* microtubules for binding neurofilaments is smaller and given by *N_MT_* − *N_on_*/*p*, resulting in a reduced on rate of

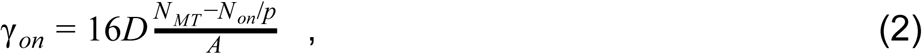

where *A* denotes the cross-sectional area of the axon.

The diffusion constant for radial neurofilament movement in axons is not known and will depend on the details of the cytoskeletal organization. Xue et al. (2015) estimated a radial diffusion coefficient for neurofilaments using the Einstein relation, i.e. *D* = *k_B_T* /*f*, which connects the diffusion coefficient *D* with the viscous drag coefficient *f*, approximated by the viscous drag coefficient of a rigid cylinder of length *l* and radius *a*, i.e. *f* = 4πμ/ ln (*l*/*a*) (Batchelor, 1970) for movement perpendicular to the axis of the cylinder. The dynamic friction coefficient μ depends on the medium the rigid cylinder moves in. Water has a dynamic friction coefficient of 1*cp* resulting in a drag coefficient of 1.1⋅10^−8^*kg*/*s* and a neurofilament diffusion coefficient of 0.36μ*m*^2^/*s* at room temperature, assuming a neurofilament radius of 20nm and a length of 5μ*m*. This diffusion coefficient results in simulated on rates that are far larger than the observed on rates, which are of the order of 10^−5^ − 10^−3^/*s*. Given the likely entanglement of neurofilaments with other structures and other neurofilaments as they move radially in the axon as well as the additional dissipation of energy by the large numbers of flexible side arms interacting with the intracellular fluid, the above calculations probably underestimate the drag coefficient. For example, a dynamic friction coefficient of 5000*cp*, as suggested for a neurofilament gel (Leterrier and Eyer, 1987), results in a drag coefficient of 5.7⋅10^−5^*kg*/*s* and a diffusion coefficient of 7.2⋅10^−5^μ*m*^2^/*s*. Because of this uncertainty, we constrain the diffusion coefficient here to produce on rates on the order of 10^−4^/*s* (Walker et al, 2019, Trivedi et al, 2007, Jung and Brown, 2009) and neurofilament transport velocities on the order of around 0.5*mm*/*day*, corresponding to the velocity of neurofilament transport in mouse ventral root and sciatic nerve motor axons in vivo (Xu and Tung, 2001). This yields a diffusion coefficient of *D* = 2 · 10^−6^μ*m*^2^/*s*.

### Predicting axon caliber

Our goal is to build a model which, given a certain number of neurofilaments and microtubules, generates an axon with the appropriate cross-sectional area. However, in addition to neurofilaments and microtubules, there are other structures in axons that occupy space (e.g. membranous organelles) and we need to include that space in our calculations. To this end, we devise a strategy that divides the space occupied by the other structures and re-allocates it to the microtubules and neurofilaments so that each of them accounts for the space covered by other structures through effective cross-sectional areas. In order to determine the effective cross-sectional areas *A_NF_* and *A_MT_* of neurofilaments and microtubules, we require that the sum of all effective cross-sectional areas of all microtubules and neurofilaments adds up to the cross-sectional area *A* of the axon, i.e.

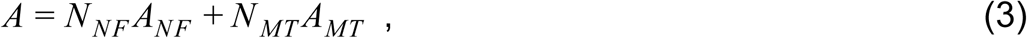

Where *N_NF_* denotes the number of neurofilaments and *N_MT_* the number of microtubules. With the densities of neurofilaments and microtubules defined by ρ*_NF_* = *N_NF_* /*A* and ρ*_MT_* = *N_MT_* /*A*, we find the relation

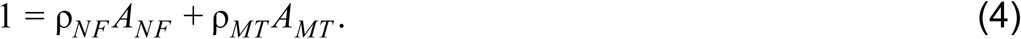

A neurofilament is composed of a backbone with a diameter of about 10*nm* and sidearms of about 15*nm* in length that are oriented perpendicular to the backbone and generate a lampbrush-like polymer structure with a diameter of about 10*nm* + 2⋅15*nm* = 40*nm*. A microtubule has an actual diameter of about 25*nm*. Using geometric considerations, we assume we can fit 5 neurofilaments around a microtubule (Lai et al, 2018), giving the fully occupied microtubule track a diameter of about 25*nm* + 2⋅40*nm* = 105*nm*. Thus, a fully occupied microtubule occupies a 6.89-fold greater cross-sectional area than a single neurofilament. We choose the ratio of the effective cross-sectional areas of neurofilaments to microtubules to reflect the ratio of the actual cross-sectional areas of these two structures, i.e. *A_MT_* = 6.89*A_NF_*, yielding, in conjunction with Eq.4, explicit values for the effective cross-sectional areas

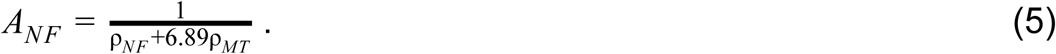

The sizes of these effective cross-sectional areas will depend on the specific type of axon being modeled and must be determined from morphometric data for the areal polymer densities in that axon type (see Figure 2B-C). For the present study, we used the data of Reles and Friede (1991) who analyzed the number of microtubules and neurofilaments with respect to axonal cross-sectional area in nodes and internodes of adult mouse sciatic nerve. A linear regression analysis of the data in Figure 5 of that study revealed microtubule and neurofilament densities of 10μ*m*^2^ and 170/μ*m*^2^ in internodes, and 53/μ*m*^2^ and 209/μ*m*^2^ in nodes of Ranvier. These densities give rise to effective cross-sectional areas of 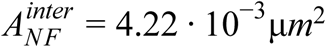 and 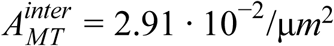 in the internodes, and 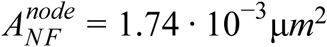 and 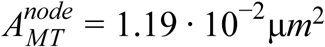 in nodes. These are the effective cross-sectional areas we use in the present study (Table 1).

**Figure 5.**
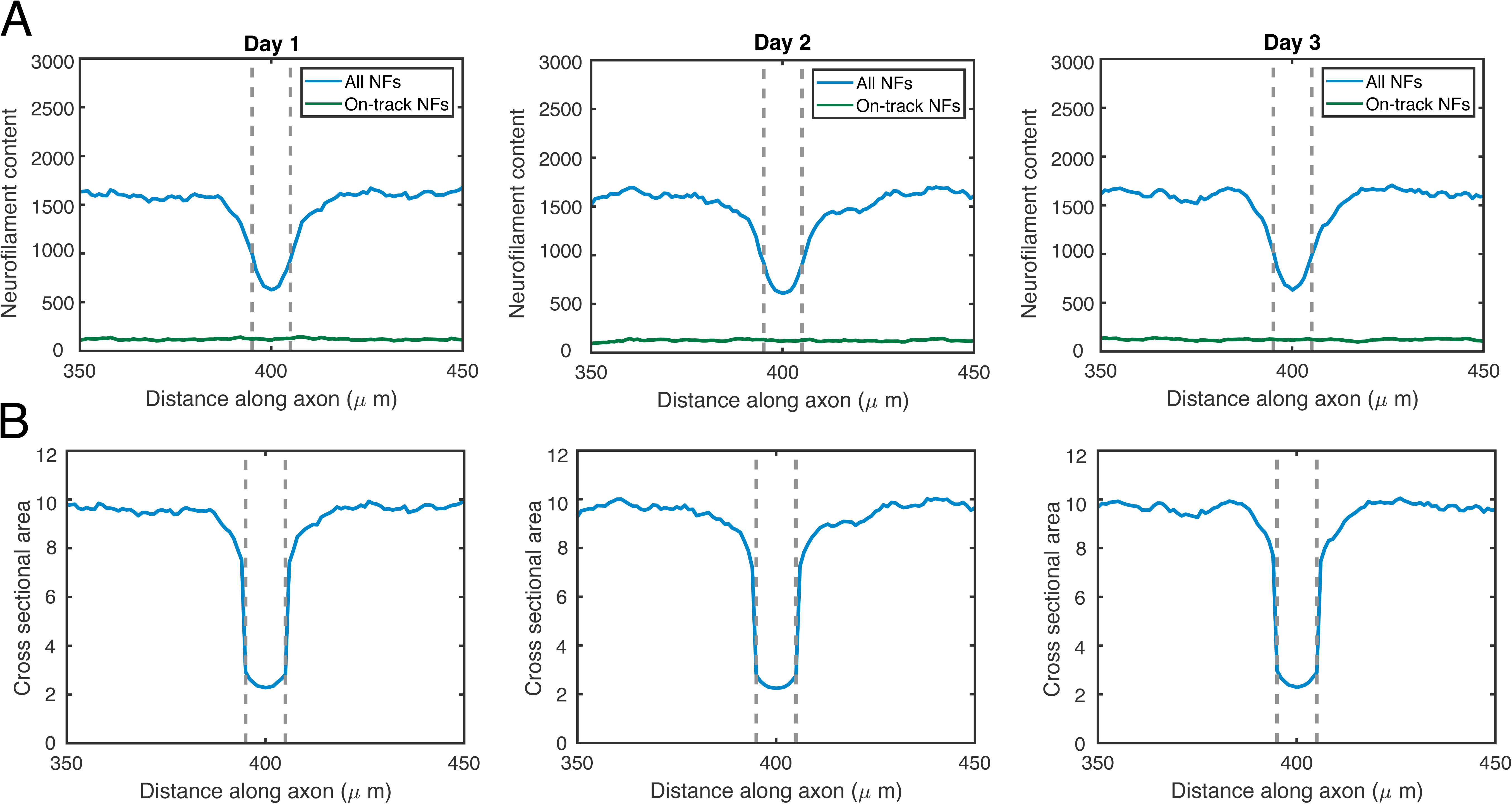
(A) Total (blue) and on-track (green) neurofilament content (neurofilament polymers per axonal cross-section) plotted versus distance along the axon at three time points during a simulation with a 10μ*m* nodal constriction. (B) Plots of the corresponding axon cross-sectional areas versus distance along the axon.

**Table 1.** Model parameters.

| Parameter | Value | Source |
| --- | --- | --- |
| Mean neurofilament length | $5.5 \mu m$ | Fenn et al, 2018 |
| Neurofilament/microtubule ratio | 16 | Reles and Friede, 1991 |
| Neurofilament density | $160/\mu m^2$ | Reles and Friede, 1991 |
| Number of neurofilament traffic lanes per microtubule $p$ | 5 | Lai et al, 2018 |
| Neurofilament anterograde velocity $v_{a,max}$ | $0.5 \mu m/s$ | Jung and Brown, 2009 |
| Neurofilament retrograde velocity $v_{r,max}$ | $-0.5 \mu m/s$ | Jung and Brown, 2009 |
| Rate from on-track pausing to moving state $\gamma_{01}$ | $0.064 s^{-1}$ | Jung and Brown, 2009 |
| Rate from on-track moving to pausing state $\gamma_{10}$ | $0.14 s^{-1}$ | Jung and Brown, 2009 |
| Reversal rate from anterograde to retrograde transport $\gamma_{ar}$ | $0.0000042 s^{-1}$ | Jung and Brown, 2009 |
| Reversal rate from retrograde to anterograde transport $\gamma_{ra}$ | $0.000014 s^{-1}$ | Jung and Brown, 2009 |
| Rate from on-track pausing to off-track state $\gamma_{off}$ | $0.0045 s^{-1}$ | Jung and Brown, 2009 |
| Rate from off-track to on-track pausing state $\gamma_{on}$ | Equation 2 | This study |
| Node length (including paranodes) | $10 \mu m$ | Walker et al, 2019 |
| Time step | $1 s$ | This study |
| Neurofilament radial diffusion coefficient | $2 \cdot 10^{-6} \mu m^2/s$ | This study |
| Effective cross-sectional areas of microtubule in internode $A_{MT}^{inter}$ | $2.92 \cdot 10^{-2} \mu m^2$ | This study |
| Effective cross-sectional areas of microtubule in node $A_{MT}^{node}$ | $1.74 \cdot 10^{-3} \mu m^2$ | This study |
| Effective cross-sectional areas of neurofilament in internode $A_{NF}^{inter}$ | $4.23 \cdot 10^{-3} \mu m^2$ | This study |
| Effective cross-sectional areas of neurofilament in node $A_{NF}^{node}$ | $1.19 \cdot 10^{-2} \mu m^2$ | This study |
| Density of microtubules in internode and node | $10/\mu m^2$, $53/\mu m^2$ | This study |
| Density of neurofilaments in internode and node | $170/\mu m^2$, $209/\mu m^2$ | This study |
| Length of photoactivation window | $5 \mu m$ | Walker et al, 2019 |

The concept of the effective cross-sectional area allows us to construct a realistic axon where the number of neurofilaments and microtubules are associated with the correct axonal caliber. It allows us to study the effects of a changing neurofilament influx and microtubule content, as seen in axons subject to radial growth, as well as to study the effects of a local change of microtubule density, e.g. the node of Ranvier, which is the subject of the present study.

### Implementation of the model

We consider a spatial domain consisting of a one-dimensional 800 μ*m* axonal segment containing *N_MT_* microtubules and *N_NF_* neurofilaments. Each microtubule is considered to extend the entire length of the axonal segment whereas the neurofilaments are shorter. To track the distribution of neurofilaments along the axon in our simulations, we discretize the axonal domain into bins of 1 μ*m* in length. The length of each neurofilament is drawn from a distribution with average length of 5.5μ*m* (minimum = 1μ*m*, maximum = 43μ*m*) obtained in a recent experimental study on cultured neurons (Fenn et al. 2018). The neurofilaments are not constrained to align with the bins so neurofilament segments may occupy part of a 1 μ*m* bin; in this case, we consider the bin to be occupied if the filament extends through at least half of the bin and to be empty if the filament extends through less than half of the bin. At each time point in the simulation, we record the number of neurofilaments *N_NF_* (*n*) in each bin *n*, as well as the number of neurofilaments that are on track in each bin *N_on_*(*n*). At the start of each simulation, we distribute the neurofilaments randomly along the axon and assign them states according to the stationary distribution of their kinetic process shown in Figure 1. However, after the model equilibrates for any given set of parameters, our results are not dependent on these initial assignments.

Axons contain a certain number of microtubules and neurofilaments packed densely in a certain volume. In myelinated axons, the neurofilaments greatly outnumber the microtubules. Thus, we consider that neurofilaments compete for a finite number of tracks and allow for the constraint that two neurofilaments cannot occupy the microtubule binding site. The effect of this is that the velocity of neurofilament transport along a microtubule track is slowed in proportion to the density of neurofilament traffic on that track. To implement these rules, we consider that on-track neurofilaments have a target velocity *v_a_*_,*max*_ if moving in the anterograde direction, and a target velocity *v_r_*_,*max*_ if moving in the retrograde direction, but we only allow these speeds to be achieved if there are sufficient unoccupied microtubule binding sites in the direction of movement. For example, consider a neurofilament moving anterogradely along the axon from left to right, and denote the 1 μ*m* bin to the right of its leading end by *r*. If all *p* binding sites on each microtubule in bin *r* are occupied (i.e., if *N_on_*(*r*) ≥ *pN_MT_*), then the velocity is set to *v_a_* = 0 and the filament will not move. Otherwise, the actual velocity *v_a_* of this neurofilament in the anterograde direction is assumed to decrease linearly with the number of other on-track filaments at location *r*, and is given by:

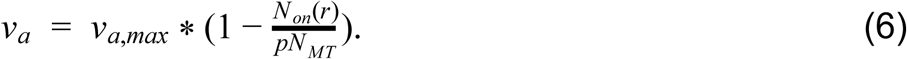

The velocity in the retrograde direction is similarly modulated by the number of on-track filaments ahead of the filament, i.e. at the bin to the left of its leading end *l*

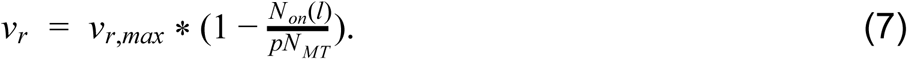

Consistent with experimental observations of neurofilament transport in cell culture (Wang et al, 2000; Wang and Brown, 2001; Uchida and Brown, 2004) and the computational modeling of neurofilament transport in vivo (Brown et al, 2005; Jung and Brown, 2009; Li et al, 2012), the reversal rates between the two directions of motion (γ*_ar_* and γ*_ra_*) are assumed to be very small (see Table 1).

Nodes of Ranvier are short spatial domains of about 1μ*m* length, where the axon is not myelinated (Figure 2). At nodes of Ranvier, the axon is constricted in area and that constriction extends both proximally and distally under the paranodal loops for a few microns. For simplicity, we refer to this as the nodal constriction, though technically it includes both the node and flanking paranodes. For the purposes of the current study, we choose 10μ*m* as the total constricted length, which is within the range encountered for large axons (Sward et al, 1995). As described above, in these constricted domains, the density of microtubules and neurofilaments is significantly larger. These changed densities result in smaller effective cross-sectional areas for the microtubules and neurofilaments. Combining Eq.2 and Eq.3, the expression for the rate γ*_on_*(*n*) at the *n*-th bin along the axon is

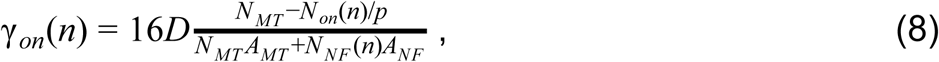

where we use the appropriate values for the effective cross-sectional areas *A_MT_* and *A_NF_* for the nodal domain and the internodal domain.

Since each neurofilament in an axonal cross-section in bin *n* experiences the same rate constants, including the rate γ*_on_*(*n*), the number of on-track neurofilaments is proportional to the total number of neurofilaments in that cross-section. As a consequence, the on rate γ*_on_* (see Eq.8), and the velocities *v_a_* and *v_r_* (Eqs.6,7) are a function of the ratio of the number of neurofilaments to microtubules. This has the consequence that the average neurofilament velocity depends only on this ratio, whereas quantities such as the flux and the cross-sectional area scale proportional to the number of neurofilaments in a cross-section.

### Simulations

The course of the simulations is summarized as follows. We choose the internodal cross-sectional area *A_inter_* of the axon we would like to consider. We use published morphometric data to determine the number of neurofilaments *N_NF_* and microtubules *N_MT_* in an internodal axonal cross-section that is associated with that size and type of axon. This determines the internodal ratio of the number of neurofilaments to microtubules, which is critical for our predictions. We then distribute neurofilaments randomly and uniformly along the axon such that the average number of neurofilaments in each cross-section equals the value *N_NF_*.

We implement the kinetic processes governing the moving and pausing neurofilaments in a stochastic manner and track the location and kinetic state of each neurofilament with time. Since a neurofilament transitions between off-track and on-track states with on rates that depend on the abundance of neurofilaments, its movement is not independent of its neighbors, i.e. the motion of one neurofilament can affect the transition rates of other neurofilaments. Thus, at each time step *dt* = 1*s* we sequentially update each neurofilament’s kinetic state and position (including the total and on-track neurofilament numbers *N_NF_* (*n*) and *N_on_*(*n*)) in all affected bins, before we proceed to the next time step.

When a neurofilament leaves the distal end of the axon it is re-inserted at the proximal end, i.e. we treat the ends of the axonal segment as a periodic boundary. Although this may seem unrealistic at first glance, it actually simulates an axonal segment in steady state where the number of neurofilaments injected per second proximally from the cell body, i.e. *j_a_*(0), is matched by the number of neurofilaments leaving per second distally away from the cell body.

The net flux of neurofilaments, i.e. the balance of anterograde and retrograde neurofilament fluxes in bin *n*, is obtained as

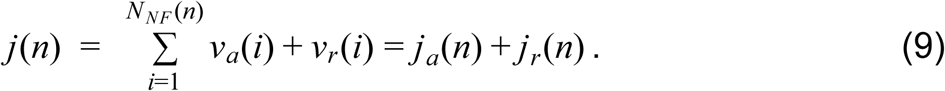

Here we sum the anterograde and retrograde velocities of each neurofilament in the moving states (given in Eqs. 6 and 7) and extending into the nth bin per bin-length, i.e. per 1μ*m*. The anterograde flux *j_a_*(*n*) is the flux that would have to be injected proximally into the axon if boundary conditions different from periodic had been used.

The ensemble-averaged velocity of all neurofilaments at a certain location (bin *n*) is obtained as the sum of velocities of all neurofilaments residing in bin *n* neurofilaments at that location

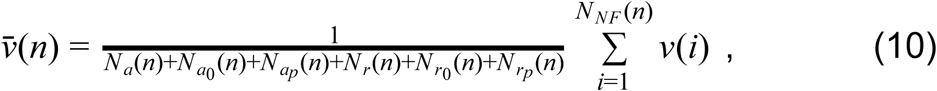

divided by the number of where *v*(*i*) denotes the velocity of neurofilament i that extends into bin *n*. Note that this velocity is determined based on the center location of neurofilament *i* as we describe in Eqs.6,7.

### Pulse-escape experiments

To simulate a fluorescence photoactivation pulse-escape experiment, we mark all segments of the neurofilaments that lie within a short window of axon of length *a* to simulate the photoactivation of neurofilaments containing PAGFP-tagged neurofilament protein. Wethen track the total length of fluorescent neurofilament polymer remaining in that activated region over time. For filaments that are partially in the activated region and partially outside of it, we only mark the segments that are within the activated region at the start of the simulation. As the photo-activated neurofilaments leave the activation window, the fluorescent intensity within the activation window declines (see Figure 10). To compare the kinetics in nodes and internodes, we choose a window at random along one of the flanking axonal internodes and compare it to a window of identical length within the node. We record the fluorescence decay at intervals of 30 seconds over 40 minutes of simulation. As in experiments, the fluorescence in the activation window at each time point is normalized to the fluorescence immediately after photoactivation and plotted against time.

**Figure 10.**
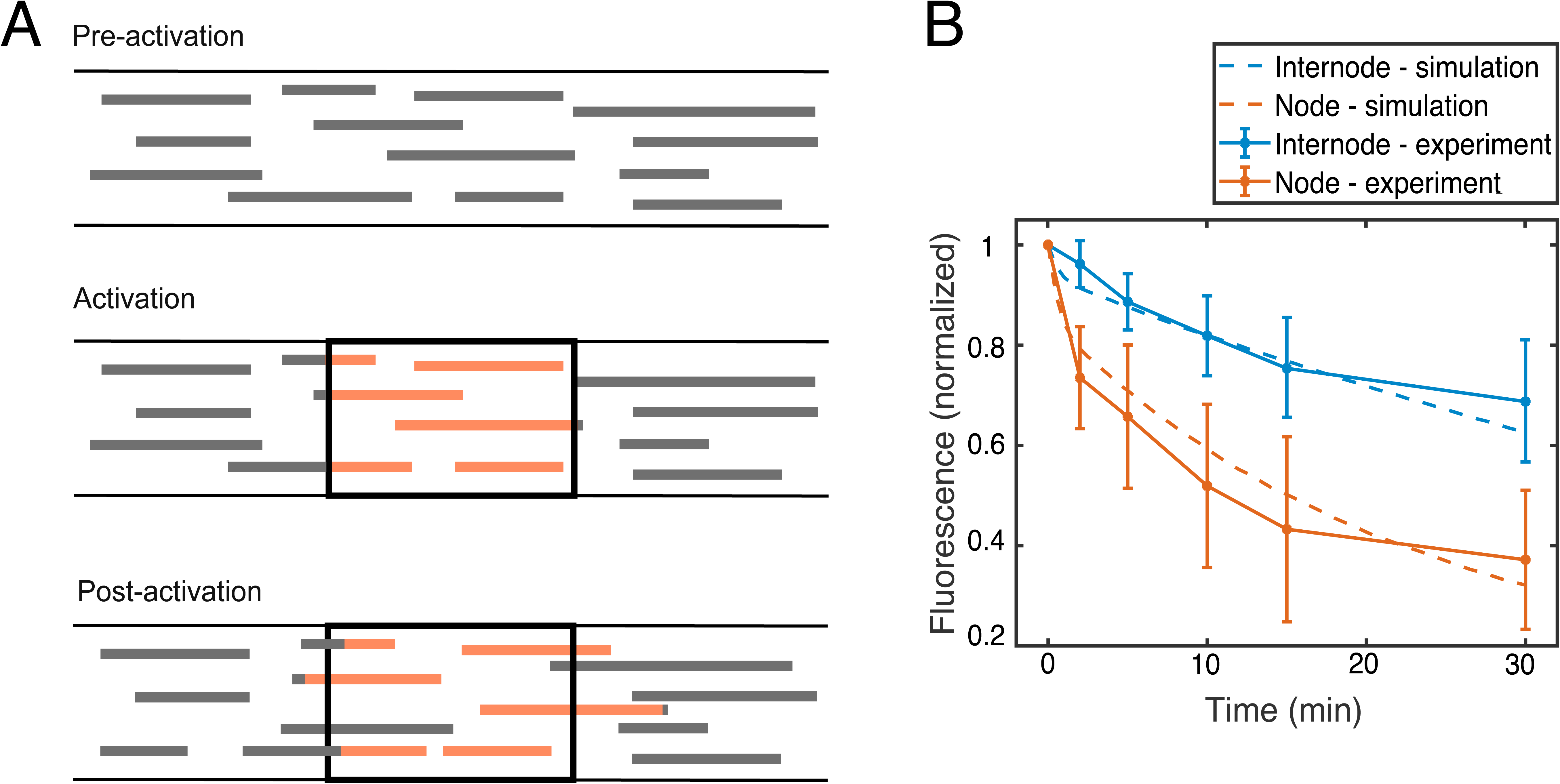
(A) Cartoon of a fluorescence photoactivation pulse-escape experiment: a population of neurofilaments within an activation window is photoactivated (red), and the fluorescence decay due to the departure of neurofilaments from the activation window is recorded over time. The decay kinetics reflect the moving and pausing behavior of the filaments (Li et al, 2014). (B) Comparison of the pulse-escape decay kinetics in our simulations (dashed lines) with our experimental data (solid lines) on contiguous nodes (orange) and internodes (blue) of myelinated axons in mouse tibial nerves (data from Walker et al, 2019). The error bars for the experimental data represent the standard deviation about the mean at each time point.

### Code availability

The simulations of neurofilament transport through model internode and node segments are run using Matlab (version R2018b). The code is available on Github (Ciocanel, 2019).

## Results

### Neurofilament transport in internodes

We choose to model axons of adult mouse sciatic nerve because there is published morphometric data on neurofilament and microtubule densities and axon caliber in nodes and internodes of these axons (Reles and Friede, 1991). To test our model, we start by simulating an internode with a cross-sectional area of 10μ*m*^2^, which corresponds to an axonal diameter of approximately 3.6μ*m*. Using the morphometric data reported in Table 1 and Fig.5 in Reles and Friede (1991), we find that such an axon will contain in cross-section about 100 microtubules and 1600 neurofilaments, corresponding to a ratio of neurofilaments to microtubules of 16. The neurofilament velocities in both anterograde and retrograde directions are modulated by the availability of open lanes along their microtubule tracks as specified in Eqs.6,7, and similarly the on rate for binding to these tracks is modulated as described in Eq.8. We initially place 100 microtubules and 213500 neurofilaments with the length distribution reported in Fenn et al. (2018) randomly along the 800 µm axonal window so that the number of neurofilaments per cross-section is about 16 times larger than the number of microtubules. The initial kinetic states of the neurofilaments are assigned randomly according to the equilibrium distribution of the kinetic states in the model, but the system rapidly equilibrates so these initial assignments do not influence the results.

Figure 3A illustrates the neurofilament distribution along the axon in contiguous 1μ*m* bins at 1, 2, and 3 days after the start of a simulation. The neurofilament content fluctuates spatially with an average standard deviation of approximately 3% about the mean. Figure 3B shows histograms of the fluctuations in neurofilament content over time, which reflects the stochastic and asynchronous movement of these cargoes. The magnitude of the fluctuations in neurofilament content about the mean is determined by the number *N_NF_* of neurofilaments in each cross-section and scales inversely proportional with 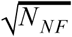 (not shown). While the stochastic fluctuations in neurofilament content along the axons are present at each point in time, there is no significant change in the distributions after 1 day. This indicates that the neurofilaments and their kinetics reached a steady state within this time. The kinetic parameters in the model dictate that the neurofilaments spend the majority of their time in off-track states, as previously validated in our experimental and computational studies (Jung and Brown, 2009; Li et al, 2012). Thus, as shown in Figure 3A, for an average number of neurofilaments per cross-section of approximately 1600, the average number of neurofilaments in the on-track states at steady state is 120 (7.5%)

To further explore the equilibration of the dynamics, we run the internode simulation for 2 hours and track average kinetic parameters of neurofilament behavior over this time interval. Figure 4 shows the mean velocity and on rate of the neurofilaments as a function of time (after initializing the simulation) during the first hours of this sample internode simulation. We obtain the mean velocities in a specified bin *i* by calculating the average of the instantaneous velocities of all neurofilaments extending over that bin. Since the internodal domain is kinetically homogeneous, we then average over all bins in the axonal window to obtain a mean velocity (and similarly, on rate) for each time. It can be seen that both the velocity and the on rate stabilize within 20 minutes of the start of the simulation. Averaging over a longer 10-hour internode simulation, we obtain a mean net velocity of 0.42 mm/day in the anterograde direction, which is in the range of published reports of 0.12-0.6 mm/day for neurofilament transport in mouse ventral root and sciatic nerve motor axons obtained using radioisotopic pulse labeling (Xu and Tung, 2001; Jung and Brown, 2009). The mean overall neurofilament on rate is estimated to be 2.52 * 10^−4^ *s*^−1^, which is similar to estimates of the on rate from fitting fluorescence photoactivation pulse-escape data in Trivedi et al, 2007; Jung and Brown, 2014. The time average (over one day) of the net neurofilament flux and the anterograde neurofilament influx (Eq.10) are obtained as ^ˉ^*j* = 7.77/*s* and ^ˉ^*j_a_* = 11.2/*s*. This means that for an axon containing 1600 neurofilaments, the net influx is predicted to be approximately 8 neurofilaments per second. The fast equilibration of the mean velocity and on rate (in less than an hour) reported in Figure 4 shows that the mean values we report are averaged over sufficiently long time intervals. It is also worth noting that these equilibrium values for velocities, fluxes, and rates are independent of the initial conditions. Our model therefore recapitulates the kinetics of neurofilament transport in internodes and can generate predictions about the flux, which have not been measured experimentally.

**Figure 3.**
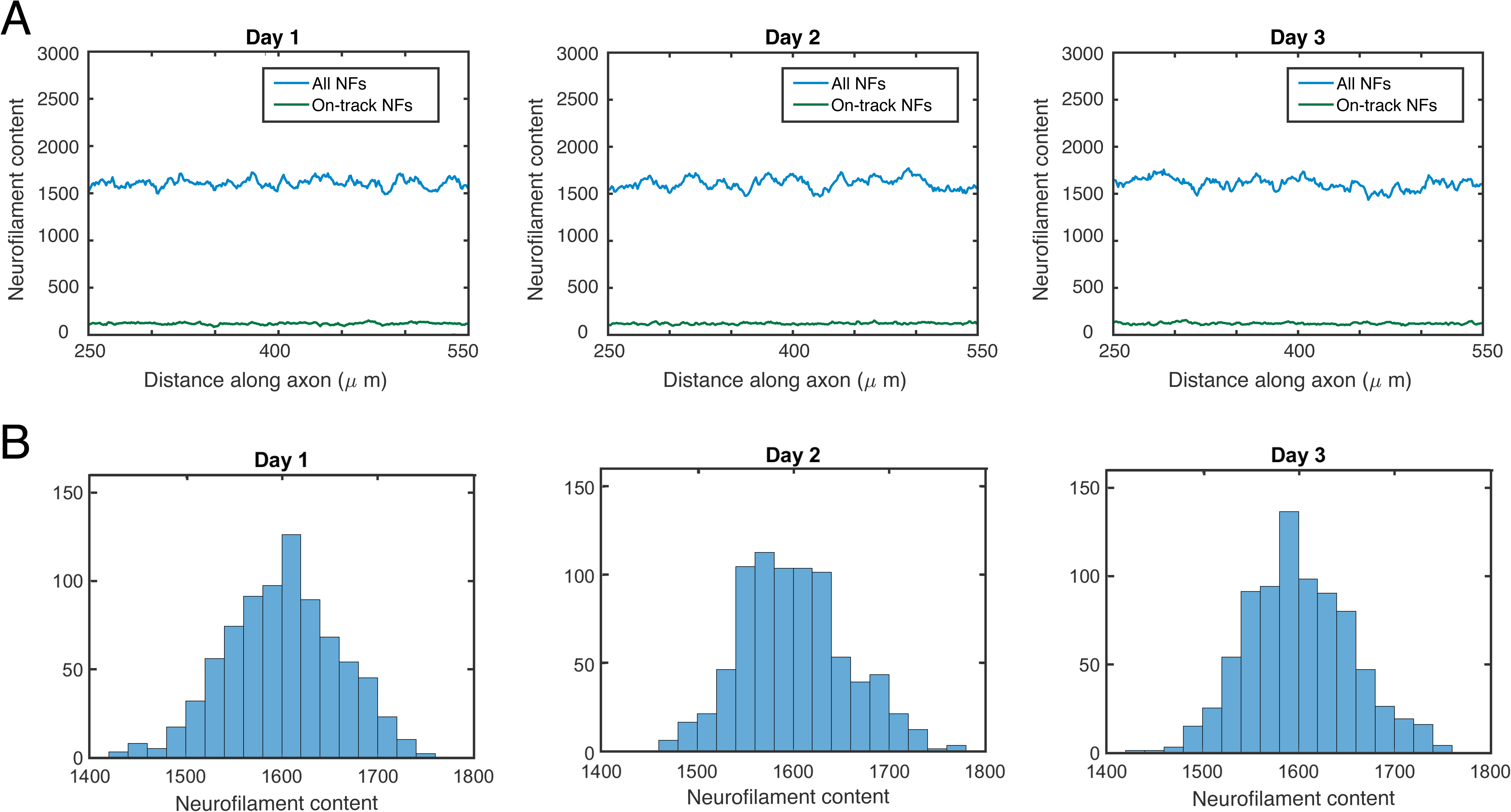
(A) Total (blue) and on-track (states *a*, *a*_0_, *r*, *r*_0_, green) neurofilament content (neurofilament number per axonal cross-section) plotted versus axon length at three time points during an internode simulation. (B) Distribution of total neurofilament content in the same internode simulation.

**Figure 4.**
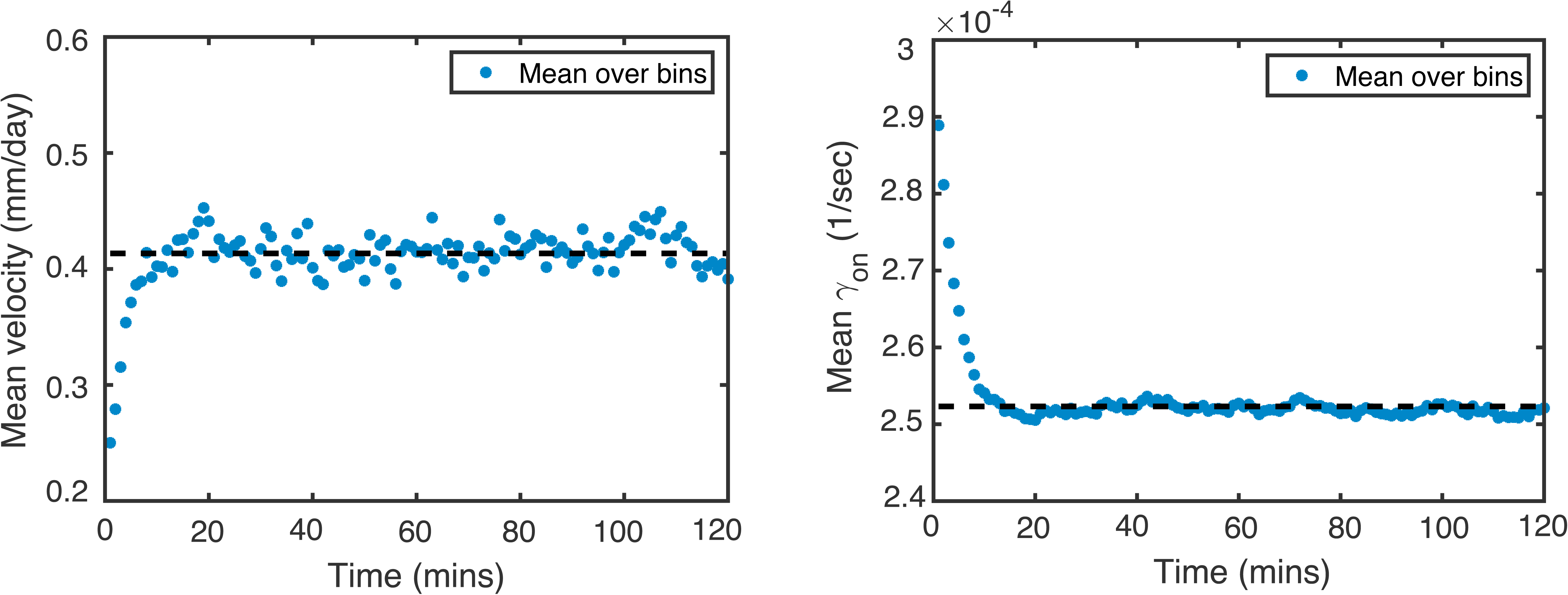
Evolution of mean velocity (left) and on rate γ*_on_* (right) with time for a simulation with no node (internode simulation). The mean velocity is calculated by averaging the velocity of all the neurofilaments within each bin, followed by averaging over all bins in the axonal domain. The dashed lines correspond to averages of mean velocities over a 10-hour period.

### Neurofilament transport across nodal constrictions

To investigate how nodes of Ranvier affect neurofilament transport, we simulate an axonal region of the same size (800μ*m*), but denote a region in the middle of this domain as a nodal constriction, dividing up the domain into two internodes. Nodes of Ranvier measure about 1μ*m* in length, but the constricted region is longer because it extends into the paranodal regions flanking the nodes. For the present study, we assume a constriction of 10μ*m* in length, which is similar to experimental measurements in large myelinated axons of adult mouse tibial nerve (Walker et al, 2019).

To determine the magnitude of the nodal constriction, we set the internodal cross-sectional area to 10μ*m*^2^ and the number of microtubules to 100 as for the internodal simulations above. Assuming that all the internodal microtubules run through the node, we then use the morphometric data in Figure 5 of Reles and Friede (1991) to extract the nodal cross-sectional area corresponding to 100 microtubules. This yields a nodal cross-sectional area of 1.76μ*m*^2^, which corresponds to a nodal constriction ratio of *A_inter_* /*A_node_* = 10/1.76 = 5.7. Using the standard deviations of the measured cross-sectional areas in the relevant axon size category provided in Table 1 in Reles and Friede (1991) (± 1.41μ*m*^2^ and ± 0.47μ*m*^2^ for the internodal and nodal cross-sectional areas, respectively) the standard deviation of the constriction ratio is estimated to be ± 1.3 (Stuart and Ord, 1994). To extract the corresponding neurofilament content ratio, i.e. the ratio of the number of neurofilaments in internodal and nodal cross-sections, we use the morphometric data in Table 1 of Reles and Friede (1991), which yields a mean ratio and standard deviation of 3.8 ± 2.3 in the relevant axon size category.

The packing densities of microtubules and neurofilaments in the node are higher than in the internode (Reles and Friede, 1991), resulting in smaller effective cross-sectional areas for these polymers (see Eq.5) (Table 1). Therefore, the nodal constriction is characterized in our simulations by a higher probability of neurofilaments engaging with microtubules, i.e. a higher on rate γ*_on_*. Since it is not known where molecular motors bind along the length of neurofilaments, we assume that the motors are uniformly distributed. This means that any bias that may be caused by motors residing proximal or distal of the center will average out. Thus, in our model we consider the motors to bind to the center of the neurofilaments.

The simulation protocol in the presence of a nodal constriction is identical to the protocol we use for the internodal simulations. We initially distribute 213500 neurofilaments uniformly along the entire 800μ*m* long axon with the same length distribution as in the internodal case, and assign the filaments to initial kinetic states randomly according to the equilibrium distribution of the kinetic states as described above. We then adjust the effective cross-sectional areas of neurofilaments and microtubules in the node as described in the Model section (see Table 1) in order to reflect the higher density of these polymers in the nodal domain. The axonal cross-sectional area is governed by the effective cross-sectional areas of these polymers in our model, so that the cross-sectional area of the node decreases. Since we define the on rate in terms of the proximity of the neurofilaments to their microtubule tracks, this higher packing density results in a larger on rate (see Eq.8) and thus neurofilaments leave this domain more rapidly than they enter. As a consequence of this imbalance, the nodal neurofilament content declines, resulting in a further decline in axonal cross-sectional area, an even larger microtubule density, and a further increase in the neurofilament velocity. This positive feedback loop results in unstable dynamics that continue until the nodal cross-sectional area and neurofilament content have declined enough that the neurofilament fluxes in the internode match those in the node. Within one day, the system attains an equilibrium resulting in a stable nodal constriction (data not shown).

In Figure 5A we show the neurofilament content across the nodes at three time points to show the stability and fluctuations in the neurofilament content and cross-sectional area once the system has reached a steady state. The colors follow the legend of Figure 3, with blue corresponding to total neurofilament content and green to on-track neurofilament content. It can be seen that there is a marked decrease in neurofilament number in the nodal constrictions (Figure 5A), which is correlated with a reduction in axonal cross-sectional area (Figure 5B). The predicted ratio of the internodal to nodal cross-sectional area (the nodal constriction ratio) is 4.3, which is slightly lower than the value of 5.7 ± 1.3 obtained from the experimental data of Reles and Friede (1991). The number of on-track neurofilaments remains constant across the node (green lines in Figure 5A), reflecting the continuity of the flow of neurofilaments. This means that the decline in neurofilament number in the node is due entirely to a decrease in the number of off-track neurofilaments. In other words, the neurofilaments move faster across the nodes by spending less time off track. This observation is consistent with findings of pulse-escape experiments in Walker et al. (2019).

It is notable that the decrease in neurofilament number is less abrupt (shallower) than the decrease in cross-sectional area. This is because the axonal cross-sectional area is determined by the effective cross-sectional areas of the polymers, which are modulated abruptly within the nodal domain, whereas the neurofilament content is determined by the neurofilament polymers which form a staggered overlapping array, with single neurofilaments often spanning the boundary between the nodal and internodal domains. We provide an animation of the nodal simulation at steady-state, with neurofilament content recorded at intervals of 30 minutes, in the Supplementary Material Video.

### Neurofilament transport velocity in nodes and internodes

We further explore neurofilament transport across the nodes of Ranvier in the above simulations by calculating average parameters of the dynamics such as velocities and on rates. At each time point, we loop through all the 1 μ*m* bins along the entire axonal domain and calculate the average of the velocities and on rates of the neurofilaments that extend into each bin. This means that for each bin, we consider not only those neurofilaments whose center is occupying that bin, but also neurofilaments in that bin that are centered in other bins. Using this approach, we obtain a mean neurofilament velocity and on rate in the internodal regions flanking the node of 0.42*mm*/*day* and 2.52 _*_ 10^−4^*s*^−1^, respectively, which are identical to the values in the earlier internode-only simulations (compare Figure 6 to Figure 4). In contrast, the mean velocity and on rate averaged across the nodal constriction are 0.93*mm*/*day* and 6.59 _*_ 10^−4^ *s*^−1^, respectively, reflecting the increased polymer density and ratio of microtubules to neurofilaments and therefore the improved access of neurofilaments to their microtubule tracks (see Figure 6B).

**Figure 6.**
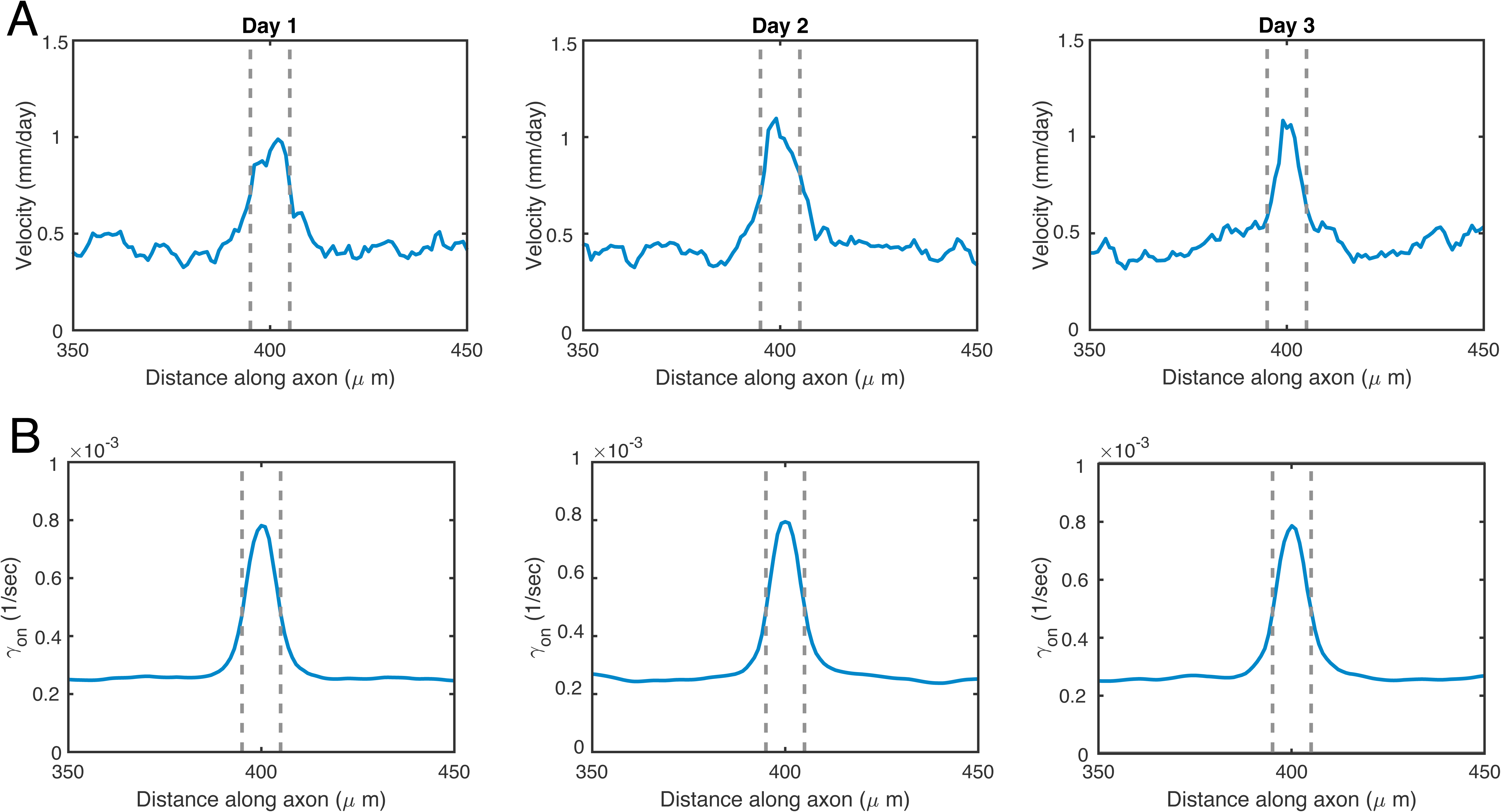
(A) Modulation of the mean velocity (governed by Eqs.6 and 7 and averaged over a three hour-window) at three time points during a simulation of neurofilament transport across a 10 μ*m* node. (B) Modulation of the corresponding mean on rates (governed by Eq.2) for the same simulations.

To quantify the effect of the nodal constriction on neurofilament content, we calculate the ratio of the mean neurofilament content in the internodes (measured as covering the 0-300 μ*m* and 500-800 μ*m* distance along the axon to avoid the node and flanking regions) versus the neurofilament content in the center (middle bin) of the node (Figure 7A). We refer to this as the content ratio. Figure 7B shows that this ratio increases over a period of several hours after the start of the simulation and then remains stable (with some fluctuations) around 2.5-3 over a period of 3 days. Note that this increase is not intended to represent how the neurofilament content ratio develops in vivo, but rather to capture the stability of the node at steady-state. This numerically predicted neurofilament content ratio is consistent with the content ratio extracted from Reles and Friede (1991), estimated to be 3.8 ± 2.3 for a 10μ*m*^2^ axon (see above). We note that the ratio of the cross-sectional areas of the internode and node (the nodal constriction ratio) shown in Figure 5B is on average 4.3 and is larger than the ratio of the number of neurofilaments in the internode and node (the neurofilament content ratio) in Figure 7B because the densities of the microtubules and neurofilaments are larger in the nodal domain.

**Figure 7.**
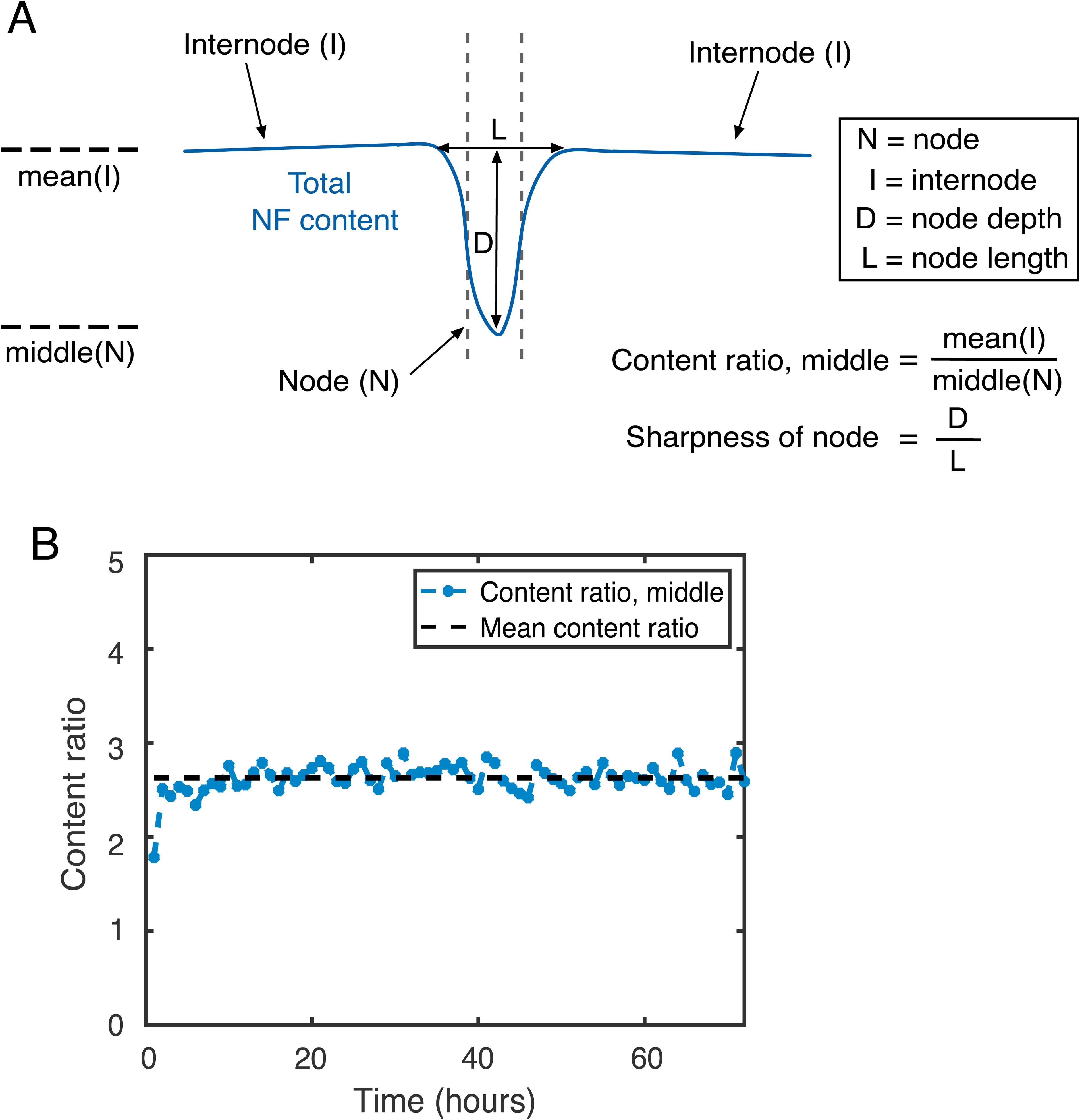
(A) Cartoon of a node simulation showing our definitions of the neurofilament content ratio (ratio between neurofilament content in the node and internode) and the “sharpness” of the nodal constriction (the depth of the node divided by its length). (B) Plot of the evolution of the neurofilament content ratio from the start of a simulation, calculated using the neurofilament content at the middle location of the node for the simulation illustrated in Figure 5A.

For the above simulations, we used a neurofilament length distribution obtained from cultured neurons (Fenn et al, 2019) because there are no published measurements of neurofilament length distribution in vivo. Since the neurofilament length distribution may be different in myelinated axons in vivo, we use our model to explore the dependence of the nodal morphometry and kinetics on neurofilament length. As for the data in cultured neurons, we assume an exponential length distribution but we vary the average length of this distribution over the range 5 *to* 25 μ*m*. We implement simulations on a 800 μ*m* domain with a 10 μ*m* node as described above. Figure 8A-B shows the neurofilament content at day 3 for each length distribution. As expected, the decrease in neurofilament content at the node is less pronounced for neurofilament lengths that exceed the length of the nodal constriction, and more pronounced at shorter neurofilament lengths. Figure 8C shows that there is a corresponding reduction in the neurofilament content ratio with increasing neurofilament length. We also explore the nodal “sharpness” by calculating the ratio D/L in Figure 8D, where D is the depth of the node and L is its length for each simulation (see cartoon in Figure 7A). Nodal sharpness also decreases as the nodes get wider and more shallow with increasing average neurofilament length. This suggests that the length distribution of neurofilaments in vivo may have significance for the neurofilament distribution across nodes, though we recognize that there are other geometric and cytoarchitectural factors that may also come into play which are beyond the scope of the current model (see Figure 2).

**Figure 8.**
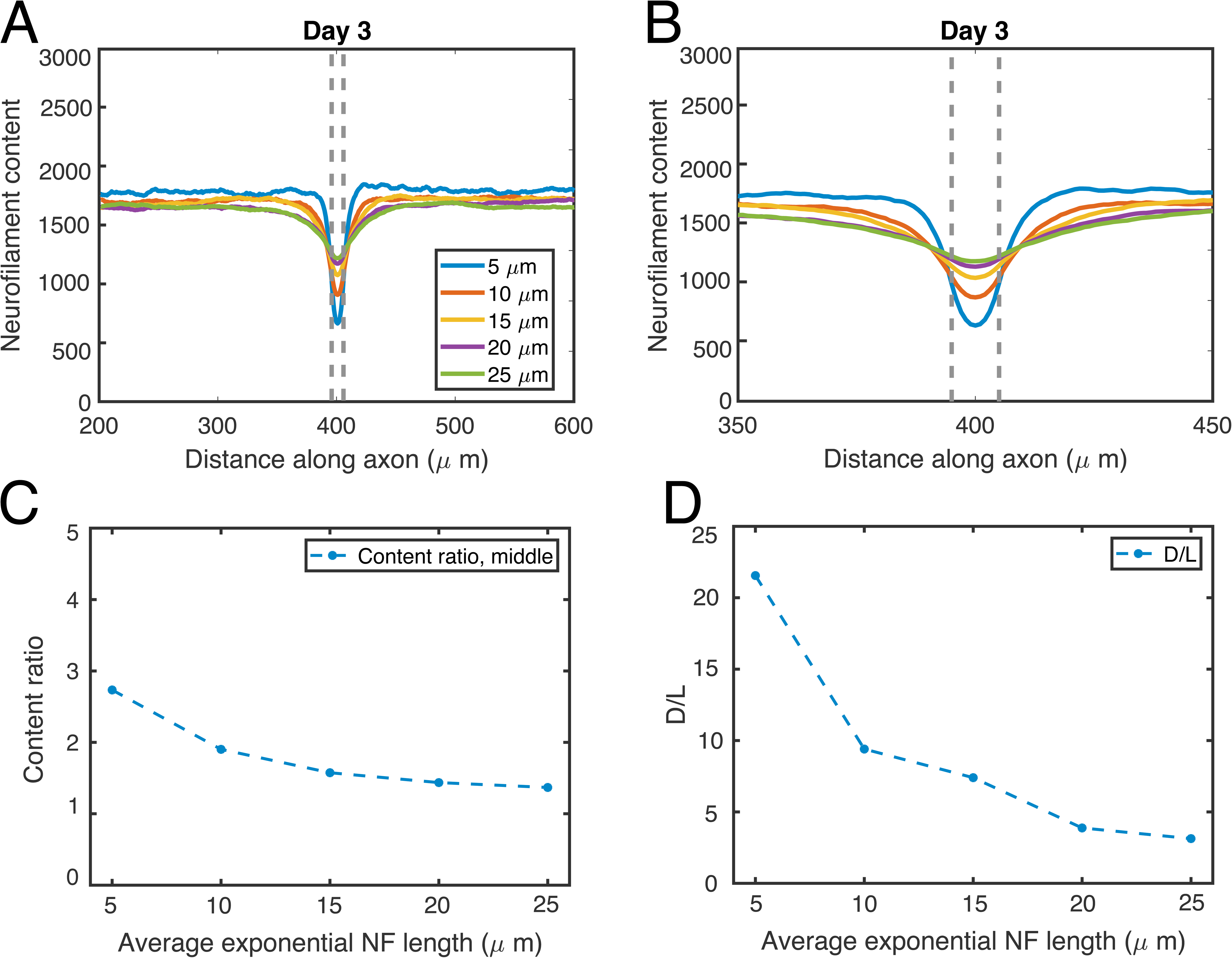
(A) Total neurofilament content at the last time point (day 3) in a node simulation, where neurofilaments are assigned lengths drawn from an exponential distribution with means ranging from 5 to 25 μ*m*. (B) Inset of (A) emphasizing the neurofilament content over a 100 μ*m* region centered at the node. (C) Dependence of the content ratio, calculated with neurofilament content at the middle of the node and averaged over 3 days of simulation, on the mean neurofilament length distribution. (D) Dependence of the fraction of node depth over node length, D/L, on the mean neurofilament length distribution.

In our simulations we considered an axon with a cross-sectional area of 10μ*m*^2^ and a ratio of neurofilaments to microtubules in the internodes of about 16. This ratio, however, can vary between axons of different calibers and axons of different types of neurons. Table-1 in Reles and Friede (1991) reports that the ratio of neurofilaments to microtubules increases with increasing internodal caliber. For example, for axons with a diameter between 3.6μ*m* and 4.2μ*m*, the ratio of the number of neurofilaments to microtubules in internodes is about 16, while for axons with a diameter smaller than 1.5μ*m*, this ratio is about 7. Similarly, Reles and Friede (1991) report that the nodal constriction ratio also increases with increasing internodal caliber, from a value of about 1.3 for axons of diameter smaller than 1.5μ*m*, to a value of 5.6 for axons with a diameter between 3.6μ*m* and 4.2μ*m*. This suggests an increase in the nodal constriction ratio with increasing ratios of neurofilaments to microtubules. Such a trend is consistent with our model predictions shown in Figure 9, which demonstrate the impact of increasing the ratio of neurofilaments to microtubules on the neurofilament distribution across the node and on the nodal neurofilament content ratio. Increasing the ratio of neurofilaments to microtubules leads to a larger decrease in neurofilament content across the nodal constriction.

**Figure 9.**
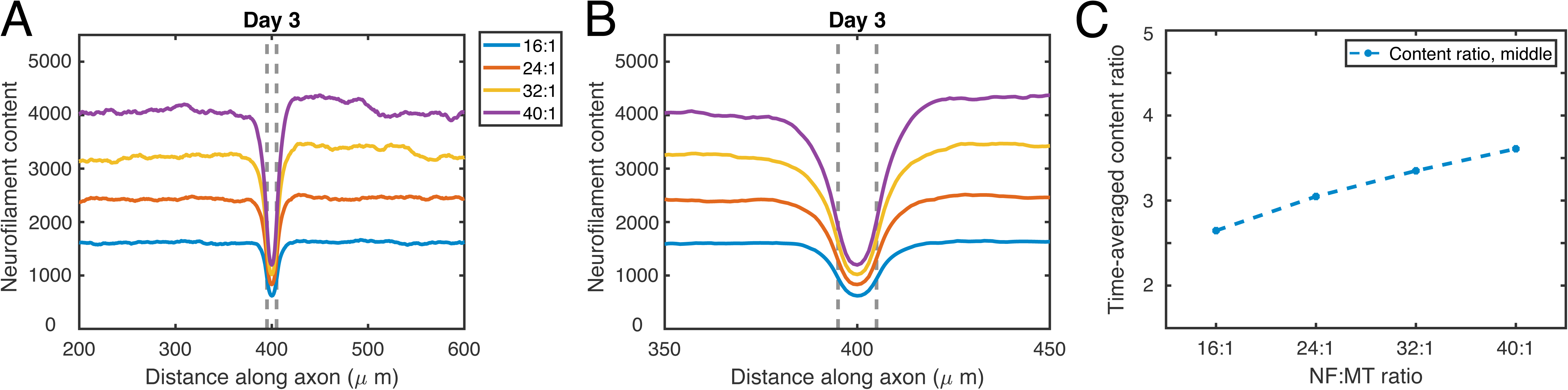
(A) Total neurofilament content at the last time point (day 3) in a node simulation, for simulations with different ratios of neurofilaments to microtubules. (B) Inset of (A) emphasizing the neurofilament content over a 100 μ*m* region centered at the node. (C) Dependence of the content ratio, calculated at the middle of the node and averaged over the last two days of the simulations, on the ratio of neurofilaments to microtubules.

### Simulated pulse-escape experiments

In our published studies on neurofilament transport in myelinated internodes in cell culture and in nodes and internodes of intact peripheral nerves ex vivo, we analyzed the transport kinetics using a fluorescence photoactivation pulse-escape technique (Monsma et al, 2014; Walker et al, 2019). In this approach, axons expressing a neurofilament protein tagged with photoactivatable green fluorescent protein (paGFP) are illuminated with violet light to activate the fluorescence in a short axonal window. Over time, fluorescent filaments depart the window by the mechanisms of axonal transport with kinetics that are dictated by the moving and pausing behavior (Figure 10A). In multiple independent studies, we have consistently observed that the decay is always biphasic, with an initial more rapidly declining phase followed by a transition to a more slowly declining phase (Trivedi et al, 2007; Alami et al, 2009, Monsma et al, 2014; Walker et al, 2019). In our model of neurofilament transport, the initial more rapidly declining phase represents the departure of neurofilaments that are on track at the time of photoactivation and thus depart within minutes. After those filaments have cleared the window, the decay transitions to a slower phase which represents the mobilization of filaments that are pausing off track and must move on track before they can depart. Thus, the initial slope of the decay curve is dictated primarily by the ratio of the on-track rate constants γ_01_ / γ_10_ later times is dictated largely by the on rate γ*_on_* in the six-state model (Figure 1) and the slope at (Li et al, 2014).

To test for overall consistency, we use the model to simulate fluorescence photoactivation pulse-escape experiments in contiguous nodes and internodes, mimicking the original experimental approach that we use to demonstrate the acceleration of nodal neurofilaments in vivo (Walker et al, 2019). Mimicking that study, we consider 10 simulated experiments with 5μ*m* activation windows and track the fluorescence decay from these windows both inside the node as well as in the adjacent internodes. Figure 10B compares the simulated and experimental data. The predicted fluorescence decay is similar to the experimental decay, and consistently fits within one standard deviation of the experimental averages. As in Walker et al (2019), we observe that the fluorescent neurofilaments leave the activated regions faster in nodes than in internodes, as evidenced by the initial slope of decay. Thus our model can explain both the published morphometric and kinetic data on neurofilament distribution and transport across axonal constrictions at nodes of Ranvier.

## Discussion

We have developed a model to test the hypothesis that the local acceleration of neurofilaments in nodes of Ranvier can be explained by a local difference in the access of these polymers to their microtubule tracks. The rationale for this hypothesis was based on two key observations in the published literature: (1) neurofilaments and microtubules are packed more densely in nodes, and (2) the ratio of neurofilaments to microtubules is lower in nodes, due largely to a local decrease in neurofilament number. Our model is based on the simple constraint that a neurofilament must be next to a microtubule in order to move along it and the observation that this is often not the case in neurofilament-rich axons, where neurofilaments greatly outnumber microtubules. Based on prior kinetic analyses, we consider that the neurofilaments alternate between distinct kinetic states which we term on-track and off-track. Neurofilaments in the on-track state move rapidly and intermittently along microtubules, pausing only briefly between bouts of movement until they disengage and become off-track. Neurofilaments in the off-track state exhibit extended pauses while they execute a radial diffusive search for another microtubule, whereupon they engage with that microtubule and move back on track. The average search time for a neurofilament to find a microtubule, which depends on the microtubule and neurofilament densities, determines the on rate. To relate axon caliber to the number of neurofilaments and microtubules, we assigned these polymers effective cross-sectional areas that we extracted from published morphometric data.

To simulate a node, we locally increased the neurofilament and microtubule density in a short segment of the axon by reducing the effective cross-sectional areas of these cytoskeletal polymers according to published measurements. This perturbation resulted in a local increase in the neurofilament on rate and a local increase in the transport velocity, yielding an emerging decline in the neurofilament content and cross-sectional area of the node, i.e. a nodal constriction. The number of filaments that were on track in the node at any point in time was similar to that in the internode (consistent with the fact that the microtubule number does not decrease in the node) but the number of filaments that were off track was reduced, resulting in an increase in the average time spent on track and thus in the average velocity, consistent with the acceleration in neurofilament transport observed experimentally in mouse tibial nerve ex vivo (Walker et al, 2019). The decrease in neurofilament number resulted in a predicted areal constriction ratio, i.e. the ratio of the internodal axonal cross-sectional area to the nodal axonal cross-sectional area, consistent with experimental observations on mouse sciatic nerve axons in vivo (Reles and Friede, 1991).

To test for consistency in our model, we compared the predicted outcomes of simulated pulse-escape experiments in the nodal and internodal domains with those observed experimentally in Walker et al. (2019). While not identical, we found that the predicted fluorescence decay in the node and internode were qualitatively similar, and within the error range of the experimental data. This is notable since the neurofilament kinetics in the node and internode in our model differ only by a single rate, the on rate, which describes differences in neurofilament access to their microtubule tracks. Thus, we conclude that the proximity of neurofilaments to microtubules is a potential regulator of neurofilament transport in axons and is sufficient to explain the local acceleration of neurofilaments through these axonal constrictions, thereby ensuring a stable morphology across these physiologically important structures.

A notable feature of our model is that it predicts an interesting interdependency between the neurofilament flux *j*, microtubule number *N_MT_*, neurofilament velocity *v*ˉ, and axonal cross-sectional area *A*. The caliber is dependent on the neurofilament content, which is determined by the neurofilament influx from the cell body and the average neurofilament velocity. The average velocity is dependent on the ratio of the number of neurofilaments and microtubules, which is influenced by the neurofilament influx. Doubling the neurofilament influx and doubling the number of microtubules results in a doubling of the cross-sectional area without any change in the ratio of neurofilaments to microtubules and thus without any change in the average neurofilament velocity. However, if the number of microtubules is held constant, even a small increase in the neurofilament flux can have a comparably large effect on the axonal cross-sectional area. For example, for the internodal parameters used in this study and 100 microtubules, increasing the influx of neurofilaments by only about 12% from 7.8/*s* to 8.7/*s* results in a decrease of the neurofilament transport velocity by 55% from 0.42*mm*/*day* to 0.19*mm*/*day* and a 2.5 times increase in the initial number of 213500 neurofilaments to 533750, leading to a doubling in the cross-sectional area from 10μ*m*^2^ to 20μ*m*^2^. This underlines the critical and in general nonlinear influence of neurofilament flux on the axon cross-sectional area: a small change in flux can result in a large change in axon caliber.

We note that our model is based on the fundamental assumption that neurofilaments perform a diffusive search to bind to available microtubule tracks. This assumption that neurofilaments can diffuse within the radial dimension of axons is consistent with prior reports that neurofilaments behave as weakly interacting elements which distribute randomly in axonal cross-sections across a range of densities (Price et al, 1988) and diffuse apart from each other freely when separated from their plasma membrane (Brown and Lasek, 1993). However, this diffusion coefficient has not been measured experimentally. Therefore, the value of this parameter in our simulations was selected by matching the resulting neurofilament on rate to within an order of magnitude of that predicted in previous work (Walker et al, 2019; Jung and Brown, 2009; Trivedi et al, 2007). Simulations with smaller diffusion coefficients and all other rate constants unchanged lead to smaller average numbers of on-track neurofilaments, as well as smaller on rates and velocities, and larger nodal constrictions (data not shown). Given the importance of the radial mobility of neurofilaments in our model, the development of methods that can measure the radial diffusion coefficient of neurofilaments in axons experimentally must be a priority for future experimentation.

Our model was designed to test the specific hypothesis at hand in a computationally efficient manner and therefore includes a number of simplifying assumptions that should be validated in future experimentation. One assumption in our model is that the neurofilament length distribution measured in cultured neurons applies to myelinated axons in vivo. We investigated the influence of this assumption in our simulations, and found that the average neurofilament length influences the sharpness of the decline in neurofilament content. The longer the neurofilaments, the more gradual the decline. To test this prediction experimentally, it will be necessary to measure the neurofilament length distribution in myelinated axons in vivo and also perform fine scale mapping of neurofilament number across nodes in these axons. A second assumption in our model is that the axon can be described as as a one-dimensional domain along which transport processes occur, incorporating the neurofilament search for microtubule tracks in the radial dimension of the axon through the proposed diffusion model. This hybrid modeling strategy has the advantage of a reduced computational cost and fewer unknown parameters. However, a more accurate approach would include the direct simulation of neurofilament motion within the crowded three-dimensional environment of the axon, incorporating the mechanical effects of the nodal morphology on neurofilaments navigating the constricted node. We plan to pursue this direction in future work, which will require experimental measurements of neurofilament polymer mechanics, organization and interactions that are currently unavailable.

Another assumption in our model is that the axonal plasma membrane deforms freely to accommodate changes in neurofilament content without resistance. A more realistic model might include a viscoelastic boundary that exhibits elastic resistance on short time scales and viscous deformation on longer time scales. However, such a model would require measurements of the deformability of the axonal plasma membrane, which will be influenced by the mechanical properties of the membrane cytoskeleton, myelin sheath and extracellular matrix. In practice, the assumption of a freely deformable boundary may be a reasonable approximation on the slow time course of axonal expansion and contraction caused by changes in neurofilament transport (i.e. hours or days). Moreover, this may not be of great consequence for our steady-state simulations of the nodes of Ranvier where the fluctuations in neurofilament content are small. In future work, where we plan to simulate the internodal axon expansion and nodal formation during development, a more detailed model of the membrane mechanics will be required.

It is important to note that nodes of Ranvier and nodal constrictions are observed in mutant mice which lack axonal neurofilaments, though both the nodes and internodes in these mice fail to attain normal caliber (Perrot et al, 2007). This is consistent with the known role of neurofilaments as space-filling structures that expand axonal caliber, but it also indicates that neurofilaments are not required for nodal constrictions to form. Thus, we do not propose that nodal constrictions arise as a consequence of the local modulation of neurofilament transport, but rather that the local modulation of neurofilament transport is essential to allow neurofilaments to navigate these constrictions without piling up on either side. In fact, nodes are complex and highly structured domains with a distinct membrane architecture that are assembled independently of neurofilaments, triggered by interactions with the myelinating cells, and likely templated by a highly organized and periodic submembrane actin cytoskeleton (Susuki et al, 2016; Ghosh et al, 2018). How the expansive forces generated by neurofilaments interact mechanically with this nodal cytoarchitecture is an intriguing question.

Neurofilaments are of clinical interest because they accumulate, sometimes excessively, in a variety of toxic neuropathies and neurodegenerative diseases, leading to swelling and distortion of axons and consequent disruption of nerve conduction (De Vos et al, 2008; Millecamps and Julien, 2013). In fact, as we have noted previously (Walker et al, 2019), it is interesting to note that studies on animal models of neurodegenerative and neurotoxic disease have often reported that these accumulations appear proximal to nodes of Ranvier (e.g., Griffin et al, 1982; Jones and Cavanagh, 1983; Jacobs, 1984; Gold et al, 1986; Hirai et al, 1999; Lancaster et al, 2018). Our model suggests that this could arise due to a partial local destabilization of axonal microtubules resulting in a reduction in microtubule number, as has been implicated in a number of neurodegenerative diseases. Thus, the requirement that neurofilaments accelerate locally in nodes to maintain a steady state morphology across these sites may make nodes particularly vulnerable to such perturbations.

## Acknowledgments

Part of this research was conducted using computational resources and services at the Ohio Supercomputer Center through the Mathematical Biosciences Institute at The Ohio State University. This project was funded in part by collaborative NSF grants IOS1656784 and IOS1656765 to AB and PJ, respectively, and NIH grant R01 NS038526 to AB. MVC was supported by The Ohio State University President’s Postdoctoral Scholars Program and by the Mathematical Biosciences Institute at The Ohio State University through NSF DMS-1440386.

## References

Alami NH, Jung P, Brown A. 2009. Myosin Va increases the efficiency of neurofilament transport by decreasing the duration of long-term pauses. Journal of Neuroscience, 29:6625–6634.

Batchelor, GK. 1970. Slender-body theory for particles of arbitrary cross-section in Stokes flow. Journal of Fluid Mechanics, 44(3): 419–440

Berthold CH. 1978. Morphology of normal peripheral axons. In Physiology and Pathobiology of Axons (edited by Waxman, SG), pp. 3–63. New York: Raven Press.

Brown A. 2014. Slow axonal transport. In “Reference Module in Biomedical Sciences”, ed. M Caplan, Elsevier

Brown A, Lasek RJ. 1993. Neurofilaments move apart freely when released from the circumferential constraint of the axonal plasma membrane. Cell Motility and the Cytoskeleton, 26(4):313–324.

Brown A, Wang L, Jung P. 2005. Stochastic simulation of neurofilament transport in axons: the “stop-and-go” hypothesis. Molecular Biology of the Cell, 16:4243–4255.

Ciocanel, MV. 2019. GitHub repository, https://github.com/veroniq04/Stochastic_neurofilament_simulation.

De Vos KJ, Grierson AJ, Ackerley S, Miller CC. 2008. Role of axonal transport in neurodegenerative diseases. Annual Review of Neuroscience, 31:151–173.

Fenn JD, Johnson CM, Peng J, Jung P, Brown A. 2018, Kymograph analysis with high temporal resolution reveals new features of neurofilament transport kinetics, Cytoskeleton (Hoboken, NJ), 75:22–41

Friede, RL, Samorajski T. 1970. Axon caliber related to neurofilaments and microtubules in sciatic nerve fibers of rats and mice. Anatomical Record, 167(4):379–87.

Ghosh A, Sherman DL, Brophy PJ. 2018. The Axonal Cytoskeleton and the Assembly of Nodes of Ranvier. The Neuroscientist, 24(2):104–110. Review.

Gold BG, Griffin JW, Price DL, Cork LC, Lowndes HE. 1986. Structural correlates of physiological abnormalities in beta, beta’-iminodipropionitrile. Brain Research, 362:205–213.

Griffin JW, Gold BG, Cork LC, Price DL, Lowndes HE. 1982. IDPN neuropathy in the cat: coexistence of proximal and distal axonal swellings. Neuropathology and Applied Neurobiology, 8:351–364.

Halter JA, Clark JJ. 1993. The influence of nodal constriction on conduction velocity in myelinated nerve fibers. Neuroreport, 4(1):89–92.

Hartline DK, Colman DR. 2007. Rapid conduction and the evolution of giant axons and myelinated fibers. Current Biology, 17(1):R29–35.

Hess A, Young JZ. 1952. The nodes of Ranvier. Proceedings of the Royal Society of London. Series B - Biological Sciences, 140(900):301–20.

Hertz P. 1909. Über den gegenseitigen durchschnittlichen Abstand von Punkten, die mit bekannter mittlerer Dichte im Raume angeordnet sind. Mathematische Annalen, 67(3):387–398

Hirai T, Mizutani M, Kimura T, Ochiai K, Umemura T, Itakura C. 1999. Neurotoxic effects of 2,5-hexanedione on normal and neurofilament-deficient quail. Toxicology and Pathology, 27:348–353.

Hoffman PN. 1995. The synthesis, axonal transport, and phosphorylation of neurofilaments determine axonal caliber in myelinated nerve fibers. The Neuroscientist, 1(2):76–83.

Hsieh ST, Kidd GJ, Crawford TO, Xu Z, Lin WM, Trapp BD, Cleveland DW, Griffin JW. 1994. Regional modulation of neurofilament organization by myelination in normal axons. Journal of Neuroscience, 4(11):6392–401.

Jacobs JM. 1984. Toxic effects on the node of ranvier. In: The Node of Ranvier (Zagoren J, ed), pp 245–272: Academic Press.

Jones HB, Cavanagh JB. 1983. Distortions of nodes of Ranvier from axonal distension by filamentous masses in hexacarbon intoxication. Journal of Neurocytology, 12:439–458.

Johnson C, Holmes WR, Brown A, Jung P. 2015. Minimizing the caliber of myelinated axons by means of nodal constrictions. Journal of Neurophysiology, 114(3):1874–84.

Jung P, Brown A. 2009. Modeling the slowing of neurofilament transport along the mouse sciatic nerve. Physical Biology, 6(4):046002.

Lancaster E, Li J, Hanania T, Liem R, Scheideler MA, Scherer SS. 2018. Myelinated axons fail to develop properly in a genetically authentic mouse model of Charcot-Marie-Tooth disease type 2E. Experimental Neurology, 308:13–25.

Leterrier JF, Eyer J. 1987. Properties of highly viscous gels formed by neurofilaments in vitro. a possible consequence of a specific inter-filament cross-bridging. Biochemical Journal, 245(1):93–101.

Li Y, Jung P, Brown A. 2012. Axonal transport of neurofilaments: a single population of intermittently moving polymers. Journal of Neuroscience, 32(2):746–58.

Li Y, Brown A, Jung P. 2014. Deciphering the axonal transport kinetics of neurofilaments using the fluorescence photo-activation pulse-escape method. BMC Neuroscience, 15(1):P132.

Lai X, Brown A, Xue C. 2018. A stochastic model that explains axonal organelle pileups induced by a reduction of molecular motors. Journal of the Royal Society Interface, 15:20180430

Millecamps S, Julien JP. 2013. Axonal transport deficits and neurodegenerative diseases. Nature Reviews Neuroscience, 14:161–176.

Monsma PC, Li Y, Fenn JD, Jung P, Brown A. 2014. Local regulation of neurofilament transport by myelinating cells. Journal of Neuroscience, 34:2979–88

Perrot R, Lonchampt P, Peterson AC, Eyer J. 2007. Axonal neurofilaments control multiple fiber properties but do not influence structure or spacing of nodes of Ranvier. Journal of Neuroscience, 27(36):9573–84.

Price RL, Paggi P, Lasek RJ, Katz MJ. 1988. Neurofilaments are spaced randomly in the radial dimension of axons. Journal of Neurocytology, 7(1):55–62.

Price RL, Lasek RJ, Katz MJ. 1990. Internal axonal cytoarchitecture is shaped locally by external compressive forces. Brain Research, 530(2):205–14.

Price RL, Lasek RJ, Katz MJ. 1993. Neurofilaments assume a less random architecture at nodes and in other regions of axonal compression. Brain Research, 607(1-2):125–33.

Reles A, Friede RL. 1991. Axonal cytoskeleton at the nodes of Ranvier. Journal of Neurocytology, 20(6):450–8.

Redner S. 2001 A Guide to First Passage Processes, Cambridge University Press, Cambridge UK.

Rydmark M. 1981. Nodal axon diameter correlates linearly with internodal axon diameter in spinal roots of the cat. Neuroscience Letters, 24(3):247–50.

Salzer, JL. 2003. Polarized domains of myelinated axons. Neuron, 40(2):297–318.

Stämpfli, R. 1954. Saltatory conduction in nerve. Physiological Reviews, 34(1):101–112.

Stuart A, Ord K. 1994. Kendall’s Advanced Theory of Statistics, Volume1: Distribution Theory,” 6th Edition, Holder Arnold, London.

Susuki K, Otani Y, Rasband MN. 2016. Submembranous cytoskeletons stabilize nodes of Ranvier. Experimental Neurology, 283(Pt B):446–51. Review.

Swärd C, Berthold CH, Nilsson-Remahl I, Rydmark M. 1995. Axonal constriction at Ranvier’s node increases during development. Neuroscience Letters, 190(3):159–62.

Trivedi N, Jung P, Brown A. 2007. Neurofilaments switch between distinct mobile and stationary states during their transport along axons. Journal of Neuroscience, 27(3):507–16.

Tsukita S, Ishikawa H. 1981. The cytoskeleton in myelinated axons: serial section study. Biomedical Research, 2(4):424–37.

Uchida A and Brown A. 2004. Arrival, reversal, and departure of neurofilaments at the tips of growing axons Molecular Biology of the Cell, 15:4215–25.

Walker CL, Uchida A, Li Y, Trivedi N, Fenn JD, Monsma PC, Lariviére RC, Julien JP, Jung P, Brown A. 2019. Local acceleration of neurofilament transport at nodes of Ranvier. Journal of Neuroscience, 39(4):663–77.

Wang L, Ho CL, Sun D, Liem RK and Brown A. 2000. Rapid movement of axonal neurofilaments interrupted by prolonged pauses. Nature Cell Biology, 2:137–41.

Wang L, Brown A. 2001. Rapid intermittent movement of axonal neurofilaments observed by fluorescence photobleaching Molecular Biology of the Cell, 12:3257–67.

Waxman SG. 1980. Determinants of conduction velocity in myelinated nerve fibers. Muscle and Nerve: Official Journal of the American Association of Electrodiagnostic Medicine, 3(2):141–50.

Xu Z, Tung VW. 2001. Temporal and spatial variations in slow axonal transport velocity along peripheral motoneuron axons. Neuroscience, 102(1):193–200.

Xue C, Shtylla B, Brown A. 2015. A stochastic multiscale model that explains the segregation of axonal microtubules and neurofilaments in neurological diseases. PLoS Computational Biology, 11(8):e1004406.

